# A thermodynamic framework for nonequilibrium self-assembly and force morphology tradeoffs in branched actin networks

**DOI:** 10.1101/2023.11.19.567734

**Authors:** Elisabeth Rennert, Suriyanarayanan Vaikuntanathan

## Abstract

Branched actin networks are involved in a variety of cellular processes, most notably the formation of lamellipodia in the leading edge of the cell. These systems adapt to varying loads through force dependent assembly rates that allow the network density and material properties to be modulated. Recent experimental work has described growth and force feedback mechanisms in these systems. Here, we consider the role played by energy dissipation in determining the kind of growth-force-morphology curves obtained in experiments. We construct a minimal model of the branched actin network self assembly process incorporating some of the established mechanisms. Our minimal analytically tractable model is able to reproduce experimental trends in density and growth rate. Further, we show how these trends depend crucially on entropy dissipation and change quantitatively if the entropy dissipation is parametrically set to values corresponding to a quasistatic state. Finally, we also identify the potential energy costs of adaptive behavior by branched actin networks, using insights from our minimal models. We suggest that the dissipative cost in the system beyond what is necessary to move the load may be necessary to maintain an adaptive steady state. Our results hence show how constraints from stochastic thermodynamics and non-equilibrium thermodynamics may bound or constrain the structures that result in such force generating processes.

## I. INTRODUCTION

The creation and transmission of forces within a cell is a crucial function of the cytoskeleton, allowing the cell to modulate its shape to in service of a variety of processes such as division, endocytosis, embryonic development, and motility through complex environments [1–4]. Branched actin networks are a component of the cytoskeleton that produces a coordinated pushing force localized to the membrane [5, 6]. This is in contrast to the pulling forces generated by kinesins and other motor proteins elsewhere in the cytoskeleton [7]. Branched networks are characterized by their branched structure where new filaments are nucleated off the side of existing ones, through a branching factor such as Arp2/3 [8]. The filaments can then polymerize and push against the load until they are capped, preventing further growth (Fig. 1). This structure gives it unique mechanical and elastic properties, distinct from crosslinked or entangled networks [5]. The assembly mechanisms of the network growing against a membrane gives rise to force dependent rates of assembly, and allows branched networks to adapt to varying mechanical forces, allowing cells to navigate through heterogeneous environments [2]. A major function of these relationships is to drive motility in cells both internally through development of lamellipodia or externally as comet tails propelling invading bacteria [9].

**Figure 1.**
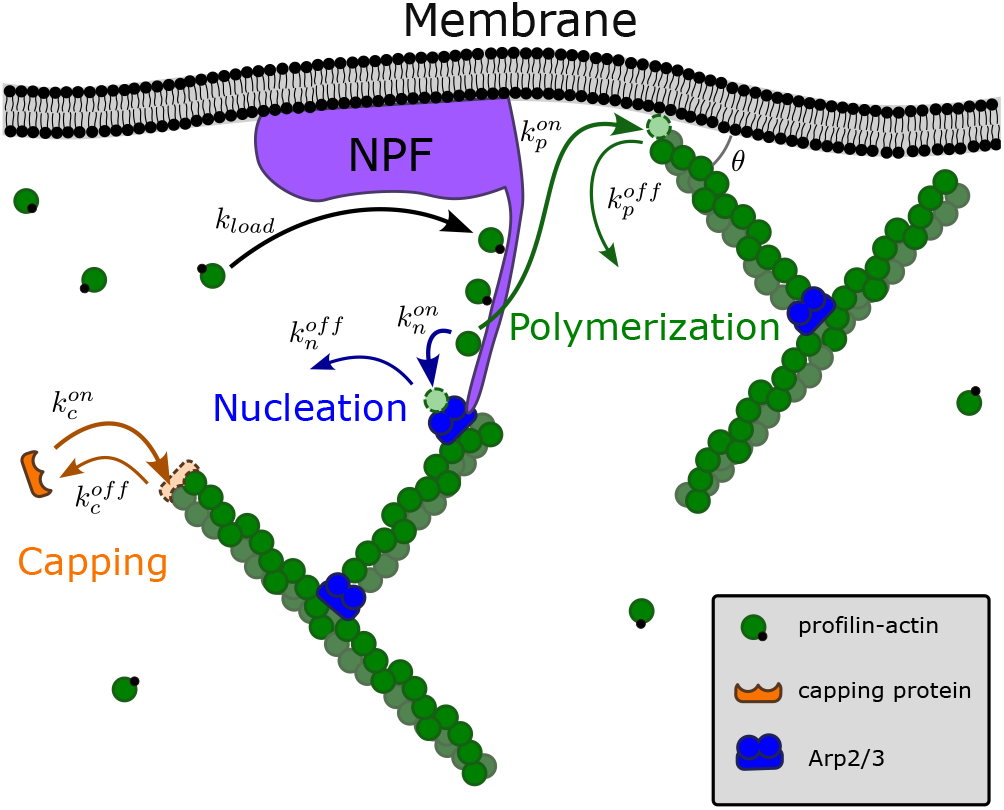
NPFs concentrate actin monomers at the membrane. From there they transfer to growing filaments or nucleate new ones. Capping protein prevents further polymerization. The Arp2/3 branching factor interacts with a region of the NPF to transfer an actin monomer to activate the branching factor. Free ends can also polymerize with actin monomers from the solution, but grow four times faster with monomers transferred from the NPF [24], ensuring that network growth remains directed toward the membrane. Free ends interact with the membrane at angle *θ*. The on and off rates describe the association and dissociation rates for each of the binding events.

There has been extensive experimental work investigating the molecular mechanisms of branched network assembly and load response [5, 10–1 However, *in vivo* these networks are difficult to isolate to probe force responses [11, 14], making it preferable to use reconstituted components *in vitro*. Additionally the filament and monomer level network structure can only be resolved with electron microscopy techniques [15, 16], so approximations from bulk network data are frequently used to describe the growing network. Reconstituted networks are frequently grown on surfaces coated in nucleation promoting factor (NPF) in an atomic force microscopy (AFM) cantilever, which provides load and displacement information relating to the network’s elastic properties and growth rate. This information can be combined with fluorescence imaging, which provides densities and incorporation rates of the various species as the network grows, to allow interrogation of the microscopic processes driving branched network growth [5, 10].

When the branched network is growing against a load, the network density and growth rate are sensitive to force. Higher load results in increased filament density, decreased growth velocity, and an increase in efficiency of the network as a molecular motor [5]. This force response is caused by changes in the network structure, both through shifts in the interaction angle of filaments with the membrane [17] and the density of growing filaments [5, 16]. This force feedback response allows the network to respond to and modulate membrane tension to create nonequilibrium shapes as well as adapt to heterogeneous extracellular environments [2, 4, 18].

The complexity generated by this limited set of interactions has motivated a variety of models and simulations. On the most complex end highly detailed whole cell simulation systems that seek to closely replicate experimental conditions [2]. However more minimal models can still capture network responses. For example, the shift in orientation angle in response to time-varying load was investigated though a Gillespie-like branch addition model [17]. An exactly solvable continuum model of an ensemble of filaments acting as pure ratchets against a fluctuating membrane with imposed inward drift can capture membrane velocity though not network density fluctuations [19]. More recently, a coarse-grained model used insights gleaned from a reinforcement learning approach to show how aspects of the external forces are encoded in the structure of the network [20]

We seek to create a minimal analytically tractable model that provides thermodynamic insight into the feedback mechanisms and force response trends in the system. Branched network assembly is a nonequilibrium process driven by ATP hydrolysis involved in actin polymerization which can serve to break detailed balance and drive the system to a nonequilibrium state even in the absence of direct force generation as seen in molecular motors. This energy input can be dissipated in maintaining the structure of the network or used to perform work moving a load, usually a membrane.

Here we show how our minimal thermodynamic model is sufficient to capture experimental trends (Fig. 6). Next, by analyzing the entropy production rate, we show that these trends are only obtainable at substantial dissipative cost. We suggest that the extra dissipative cost, beyond what might be necessary to simply push a load, might be due to the need to maintain an adaptive steady state. Indeed, while the fluctuations in network free end density dissipate energy, they may also allow the system to maintain the motion of the membrane across shifts in load as part of an Energy Speed Accuracy (ESA) tradeoff [21]. The force response has been experimentally investigated with a variety of forcing conditions (e.g. constant stiffness, oscillating load) [10, 14] which can vary the tradeoffs in the system. We will focus on the quasistatic case where the load is held constant for a given network. We note previous work has investigated thermodynamic relationships between driving forces and resulting configurations in polymer-like self-assembly systems [22, 23]. This previous work explored simpler systems, but we expand the framework to describe the dynamics of branched network assembly.

The outline of the rest of the paper is as follows. We first describe the interactions involved in assembling the physical system of branched actin networks, then introduce our minimal model. We apply approximations to self-consistently solve for the steady state. We show how our minimal thermodynamic model is consistent with experimental trends. We then calculate the entropy production rate of the growing system, and show how it can provide a parametric bound on the available configurations.

## II. OVERVIEW OF BRANCHED ACTIN NETWORKS

While there can be many components involved, branched networks can be reconstituted with only Arp2/3 branching factor, a Nucleation Promoting Factor (NPF) from the WASP family proteins, capping protein (CP), monomeric actin-profilin complex, and filamentous actin [5, 10, 25]. The NPF binds the profilin-actin complex, concentrating it at the membrane. The actin can then be transferred to the end of an existing filament and continue its polymerization or bind an Arp2/3 complex and nucleate a new filament, as described in Fig. 1. The CP binds to free ends and prevents further polymerization, meaning the filament will fall away from the growing front of the network. The growing ends of actin filaments can also bind to the NPFs which prevents nucleation and tethers the free end to the membrane in a process called barbed end interference [26]. This effect, along with competition between the Arp2/3 and the growing filaments for the pool of NPF-associated actin monomers [24] contributes to the negative feedback of uncapped free ends on the nucleation rates of new free ends [10]. Capping is necessary in the system to reduce the number of free ends in the system, which in turn increases nucleation rates as capped ends cannot bind NPFs or use NPF-associated actin monomers to polymerize [27]. Because Apr2/3 must bind to an existing filament to nucleate a new one, the nucleation process is autocatalytic. However, the negative feedback from high numbers of free ends prevent exponential nucleation [24] and allows the system to fluctuate around some average steady state network density. This steady state density depends on the rates of the various processes in the system, and can shift in response to load as the rates are impacted by Brownian Ratchet effects.

### A. Brownian Ratchet

Both polymerization and capping of filaments in contact with the membrane can be described as a Brownian Ratchet process [10, 28]. Brownian ratchets are a theoretical framework to describe force generation through thermal fluctuations and steric hindrance. When a filament is in contact with the membrane, a new monomer cannot be added unless fluctuations create a large enough gap to intercalate it. For a filament growing at angle *θ* to the membrane, the gap size would be Δ = *δ* sin *θ*, where *δ* is the monomer size. To move a load *f* that distance requires energy *C*_*f*_ = *f* Δ, so the probability it will occur though thermal fluctuations is would be proportional to exp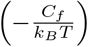 [29]. For an ensemble of filaments polymerizing in contact with the membrane, we assume the load is evenly distributed across all free ends (*E*), scaling the polymerization rate by exp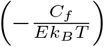.

The average free end length of a filament in the network is determined by the ratio of polymerization and capping rates. Because both these processes are Brownian ratchets, the rates scale exponentially with the energy required to move the membrane. Both processes occur at the same site on a free end, so the difference in energy cost to add each respective monomer type depends only on the size, *δ*. Both monomers are similar in size, allowing the rates to scale together with load, and keeping the ratio of the rates and thus the free end lengths constant, as has been observed experimentally [5]. The Brownian Ratchet component of the capping rate is crucial to the force response feedback loop in the network. If the number of free ends is higher than the steady state value, capping is amplified due to lower per-filament load, which in turn reduces E. Conversely, if the density is low, polymerization and capping are both suppressed and nucleation is encouraged, allowing the system to return to the steady state average.

## III. A MINIMAL THERMODYNAMIC MODEL FOR BRANCHED ACTIN NETWORK GROWTH AND ASSEMBLY

To investigate the thermodynamic bounds on the structural fluctuations in this type of system, we develop a minimal model that still captures the force response trends observed in experiments. This model emulates a column of branched actin network growing against a load, as is frequently employed in experimental setups [5, 10, 12, 14].

As our model assembles, it is characterized by the number of free ends (E), which is analogous to the network density and impacts the rates at which components are incorporated. We assume spatial homogeneity as well as a constant load, so the load is evenly distributed across all free ends and the per-filament load is determined by E, allowing us to model the system as a one dimensional series of nucleation (N), polymerization (P), and capping events (C), that can be thought of like monomers added to a growing chain. We further assume a large enough reservoir that background monomer concentrations remain constant as the network grows. This allows us to define constant per-filament addition rates for the zero load case, where the Brownian Ratchet component is 1. These rates are based on the background monomer concentrations and the 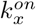 association rates, where *x* ∈ {*n, p, c*}, as shown in Fig. 1,

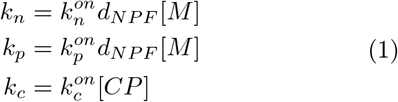

where *d*_*NPF*_ is the density of NPFs at the load, [M] is the concentration of soluble actin monomers, and [CP] is the concentration of capping protein. The corresponding removal rates are simply the 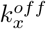 rates.

Nucleation and capping events respectively increase or decrease, the number of free ends (E), while polymerization moves the load forward. The addition and removal rates for the events depend on the overall configuration of the system are depicted in Fig. 2, with only the most recent event able to be removed, in which case the second most recent is now the ‘tip’ of the chain. This linear picture is meant to approximate tracing a single path or ‘filament of interest’ through an existing network and can be viewed as a mean field model. We then take that sequence of events to be representative of the average structure of the network as a whole.

**Figure 2.**
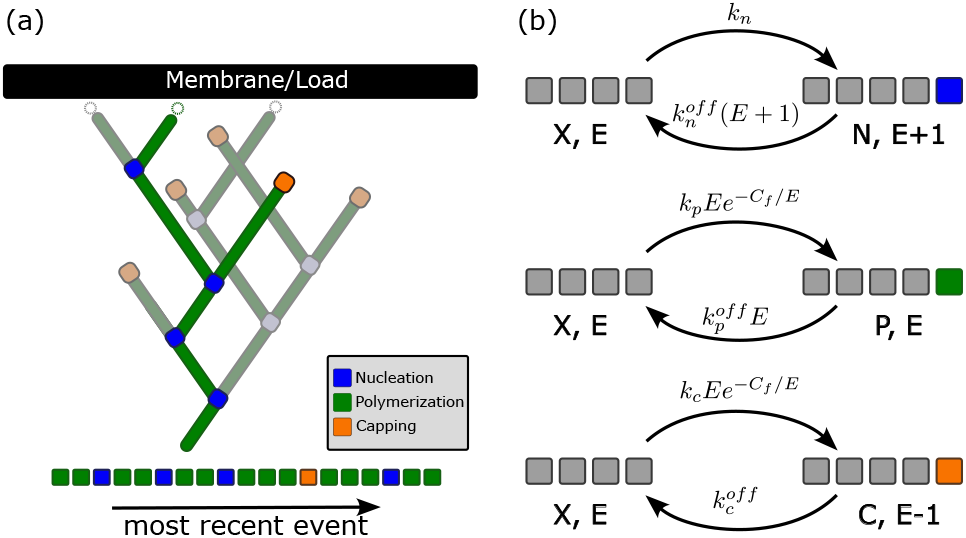
Depiction of full model. (a) Representation of a state in the full model. The full order of events that have been incorporated into the chain is recorded, emulating a path traced through a full branched network growing. Desaturated components are what is approximated to be the structure of the complete network based on the composition of the filament of interest, resulting in some number of active free ends at the membrane. (b) Transition rates in the full model. *X* ∈ {*N, P, C*} Grey squares can represent any type of event, as addition and removal rates depend only on the current number of free ends and most recent event.

To make analytical progress we make one further approximation (which is justified subsequently). In the model depicted in Fig. 2, the history of the events on the filament of interest is completely captured as the order of events is preserved. However, this model’s infinite state space make it difficult to solve the master equation, so we will instead consider an effective Markov state model defined by the most recent event and the number of free ends, *E*, shown in Fig. S1. In this simplified effective model, the absence of a history results in uncertainty about the appropriate state to transition to upon event removal. Prediction of the next state relies on calculating conditional probabilities based on the second most recent event. In the case described in Fig. 2B top, the reverse rate for the effective model would become 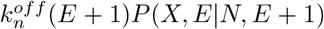. Because the number of free ends is prescribed by the type of the removed event the conditional probability can be expressed simply as *P*(*X*|*E*).

Due to the autocatalytic nature of filament nucleation with the negative feedback of *E*, the number of free ends in a growing network fluctuates around some steady state *E*_*ss*_ = ⟨*E*⟩, representing the most probable value of *E*. Our objective is to replicate the growth and force response of a network comprising numerous polymerizing filaments in a minimal model that can be analyzed thermodynamically. Given that these are bulk properties resulting from an ensemble of filaments, we prioritize the average behavior over extreme and infrequent states. To capture the bulk steady state behavior of the network the we can approximate

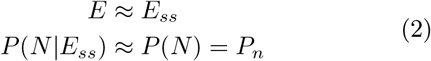

when *E* is close to *E*_*ss*_, which is where the system spends the most time.

### A. An analytical steady state solution

A consequence of our steady state assumption is that the state probabilities are proportional to the flux of that event type into the system *J*_*x*_. This means that

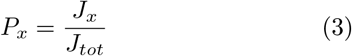

Where *J*_*tot*_ = *J*_*n*_ + *J*_*p*_ + *J*_*c*_. For there to be a steady state *E*, the flux of nucleation and capping events into the system must be equal. So, by setting *J*_*n*_ = *J*_*c*_ we get

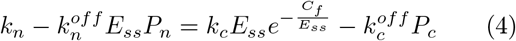

and solve for *E*_*ss*_

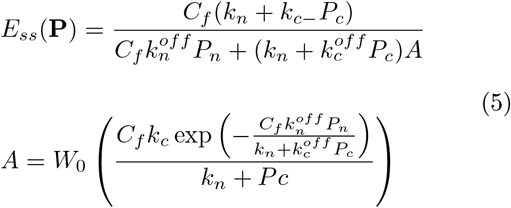

where *W*_0_(*x*) is the Lambert W function.

Eq. 5 implies that the steady state structure can be completely described by the state probabilities **P**. To solve for the steady state where 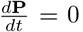, we construct a matrix **M** such that 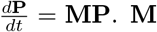 is determined by the master equation for our minimal model described in Eq. S1. The eigenvector of **M** corresponding to the zero eigenvalue will give us the steady state **P**.

Now that we have developed and solved our minimal model of branched network assembly, we verify the model through simulations and comparison to experimental findings. Once the association and dissociation rates in the system are set, we can simulate the growing system using a kinetic Monte Carlo algorithm. Events (adding a new monomer through nucleation, polymerization or capping) are successively added and removed to the chain, with rates determined by the current number of free ends, as described in the full model in Fig. 2.

The previous event is part of the current state, so conditional probabilities are not needed to identify the new tip of the the chain upon a removal event. Nucleation and capping events increment or decrement E, respectively. We evolve the system until it reaches a steady state. Once at the steady state we let the system assemble until the chain of events reaches a large enough end length to estimate the steady state event probabilities. These are calculated simply from the composition of the chain. Regardless of the initial number of free ends, the system will reach and fluctuate around a steady state *E*_*ss*_ = ⟨*E*⟩, as well as steady state event probabilities **P**.

### B. Steady state and dynamical behavior regime in various parameter regimes qualitatively resembles some experimental trends

The structures generated by the simulation using the full model allow us to calculate the probability of incorporating each event type. We then use this to self-consistently calculate the steady state solution in the effective 3-state model. The distributions agree between the simulation and the steady state solution as shown in Fig. 3B. Our results also agree with the general force response trends observed experimentally of networks growing under step-wise increasing load, with convex decay in growth velocity and increase to saturation of free ends with increasing load [5, 10]. We also investigate the behavior of spring like loads, capturing an initial plateau followed by decay of the load velocity, as shown in Fig. 4. We simulated the full model with a Hookean spring like load defined as *f* = *k*_*spring*_*x*(*t*), where *x*(*t*) is the instantaneous position of the front, and the energy to move the load one monomer length remains *C*_*f*_ = *f*Δ. The load response of this system demonstrated some of the plateau structure in seen in experimental works [12, 13]. Interestingly, higher spring constants lead to larger plateaus, likely due to slower initial velocity allowing the system to adapt more fully before reaching the high load regime where system composition begins to change.

**Figure 3.**
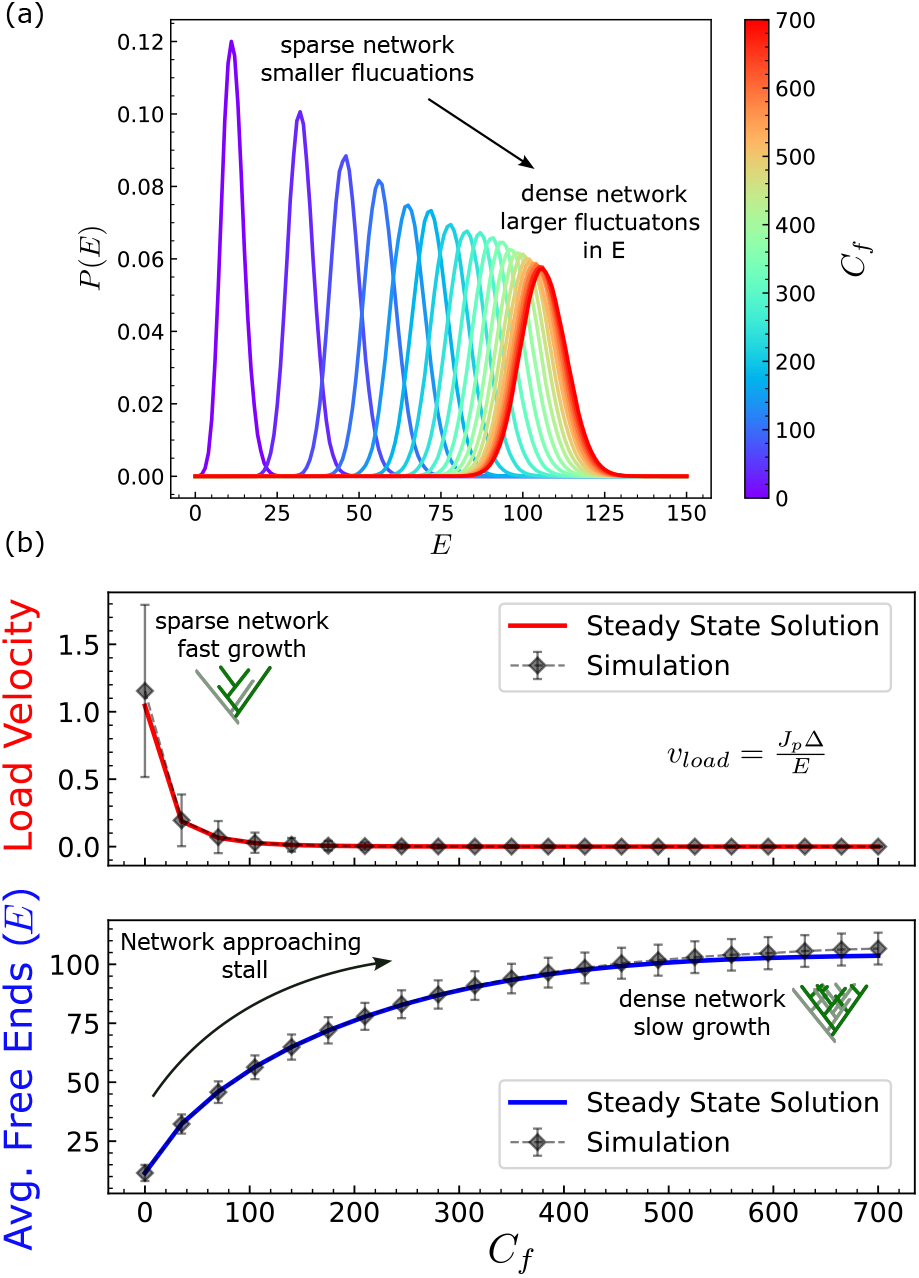
The force response of the network impacts both the steady state average and the level of fluctuations around that steady state. (a) The distribution of the number of free ends is symmetrically distributed around the steady state average. As the total network density increases and the polymerization rate slows, the increasing width of the free ends distributions indicates greater fluctuations and a fractionally lower effect of the change of the single discrete nucleation or capping event given the higher overall number of free ends. (b) There is good agreement between the simulated *E* values, and the self consistent steady state solution. The trends of free ends *E* and load velocity 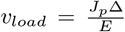, with increasing load are similar to experimental ones. The divergence of the simulated and analytic solution at high loads reflects the increasingly wide distribution of free ends with increasing load as seen in (a). However this high load state is very close to the stall load as demonstrated by the vanishingly small load velocity in these cases, and might be considered under a different regime.

**Figure 4.**
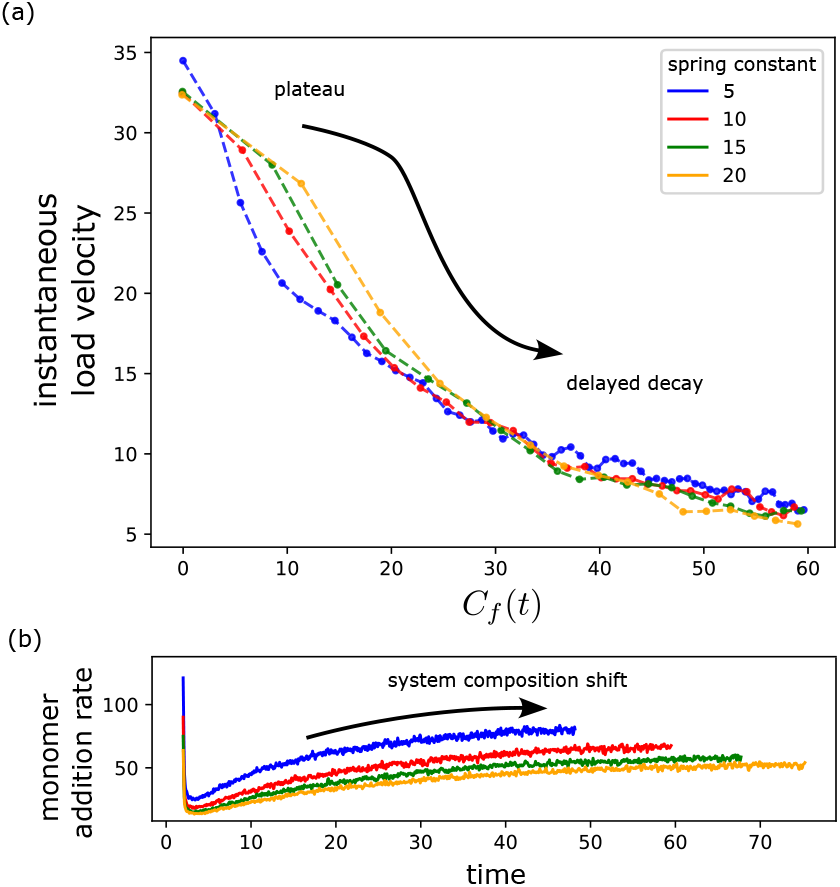
(a) The instantaneous force response of a network grown against a load with constant stiffness. In this case the load exerts a force *f* (*t*) = *x*(*t*)*k*_*spring*_, where *x*(*t*) is the current displacement of the membrane. The energy required to move the membrane on monomer gap size remains *C*_*f*_ (*t*) = *f* (*t*)Δ. This system cannot be solved for a non equilibrium steady state due to the constant change in *C*_*f*_ until the network reaches stall. However the response with an initial plateau at low *C*_*f*_ followed by decay of load velocity shows trends similar to experimental findings in cases with spring-like loads. (b) After the initial drop in monomer addition rate, it recovers as the system begins to approach stall and shift the system composition. [12, 13]

We examined a range of values of *k*_*x*_ to identify the quantities crucial to system behavior. Fig. 5 (a) [top] shows when the load velocity profiles are normalized by their c value, they seem to converge asymptotically as the *c* ≡ ∑ *k*_*x*_ is increased. Further, these results also predict that as *c* is increased, the system shifts to a polymerization driven regime (as indicated by a high value of *P*_*p*_. Hence, for the low load regime where *k*_*x*_ >> *k*_*rx*_, which is where many experimental systems operate, we may expect the force response to scale with the unloaded forward rates. This echoes experimental data in which force velocity curves are normalized by unloaded growth rates [10, 12, 13].

**Figure 5.**
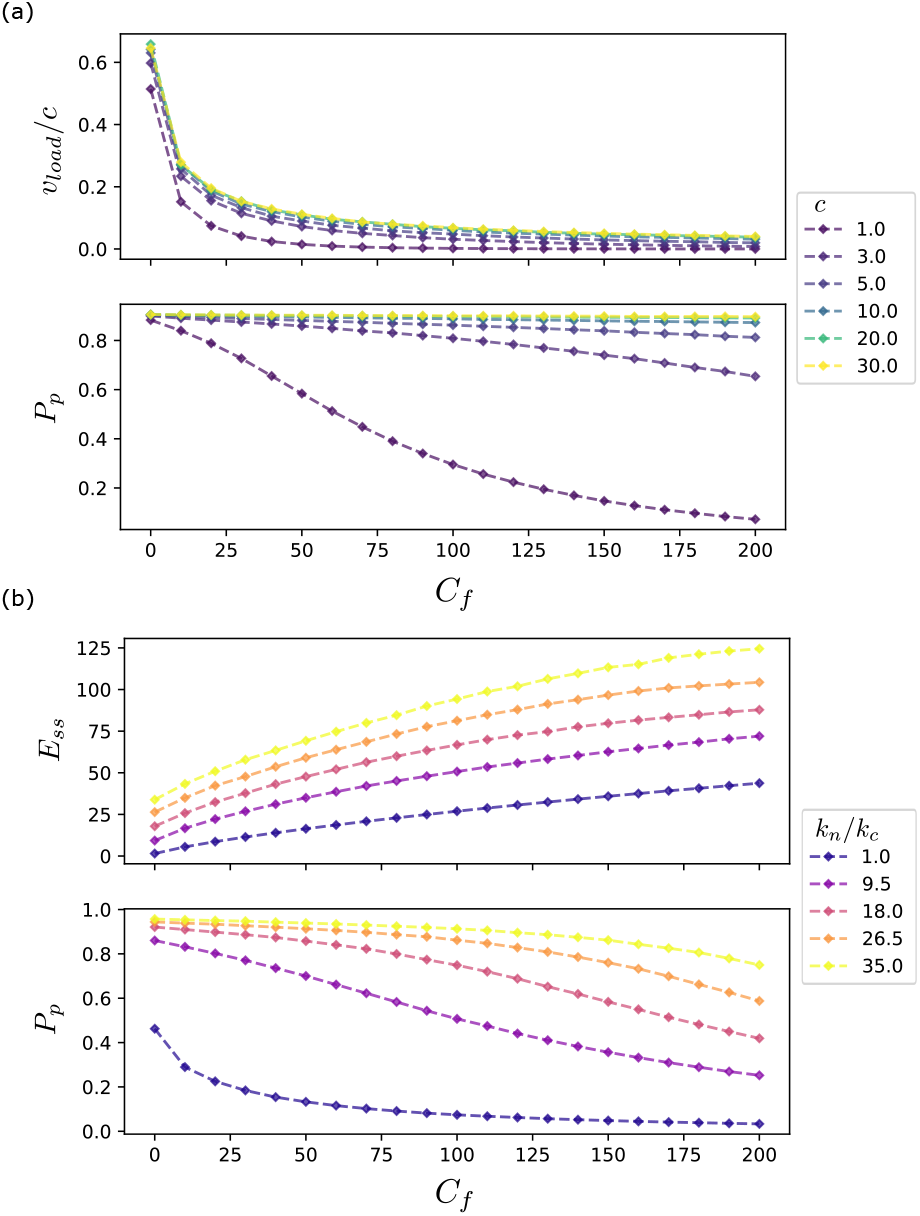
Steady state behavior in various parameter regimes of *c* = ∑*k*_*x*_ and *k*_*n*_*/k*_*c*_. (a) Load velocity profiles normalized by *c* (top) and steady state polymerization probability (bottom) for various values of *c* (b) Steady state free ends (top) and polymerization probability (bottom) profiles for various values of *k*_*n*_*/k*_*c*_

We also considered changing the ratio *k*_*n*_*/k*_*c*_ Fig. 5 (b). For a fixed *k*_*n*_*/k*_*c*_, we find that increasing the load shifts slowly the system from a polymerization dominated regime Fig. 5 (b) (bottom). This is consistent with Fig. 5 (a)(bottom). Further, the number of free ends increases with the load in a manner that is a qualitatively consistent across a range of values of *k*_*n*_*/k*_*c*_. The qualitatively consistency in our parameter scans shows how the results from our minimal model can be extended to different experimentally important regimes.

### C. Entropy production rate and thermodynamic constraints on force-morphology curves

Using the forward and reverse rates in the effective model, we define the total entropy production rate 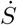, which is the sum over the entropy production of each of the possible transitions. This allows us to calculate

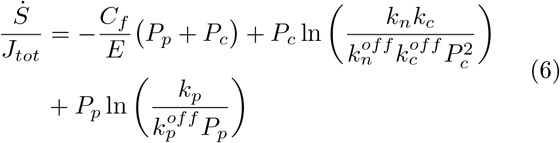

the average entropy production per monomer addition. The steady state assumption from Eq. 2 we used to simplify the model introduces some error. If *E* ≈ *E*_*ss*_ then *P* (*E*) ≈ *P* (*E*_*ss*_) ≈ 1, which would be represented by a delta function. However, the distribution *P* (*E*) in the full model, as shown in Fig. 3A, is Gaussian, with broader distributions seen at higher loads. The broader the underlying distribution of *E* values is, the farther from a delta function and thus the worse our approximation is. The increasing deviation of the distributions correlates with the slight disagreement between steady state solution and simulation seen in Fig. 3B at high load. These broader distributions are also caused by a suppression of polymerization at higher loads, so more time is spent adding and removing free ends which drives the fluctuations.

Eq. 6 provides a basis to obtain thermodynamic bounds between various quantities. For example we define

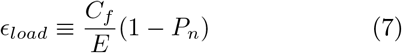

as the thermodynamic cost per monomer addition due to the load. The requirement that 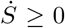 in Eq. 6 can now be used to bound and connect such thermodynamic costs to quantities that parameterize the structure such as *E* the number of free ends in the system. In Fig. 6a, we show how the non-negativity of the entropy production rate can be parametrically expressed as a bound on available states for a given load or monomer addition cost. In Fig. 6b, we describe a similar plot but now connecting the number of free ends *E* to the force on the membrane, *C*_*f*_. Specifically, the second law bound on *ϵ*_*load*_ shown in Fig. 6a can be expressed as

**Figure 6.**
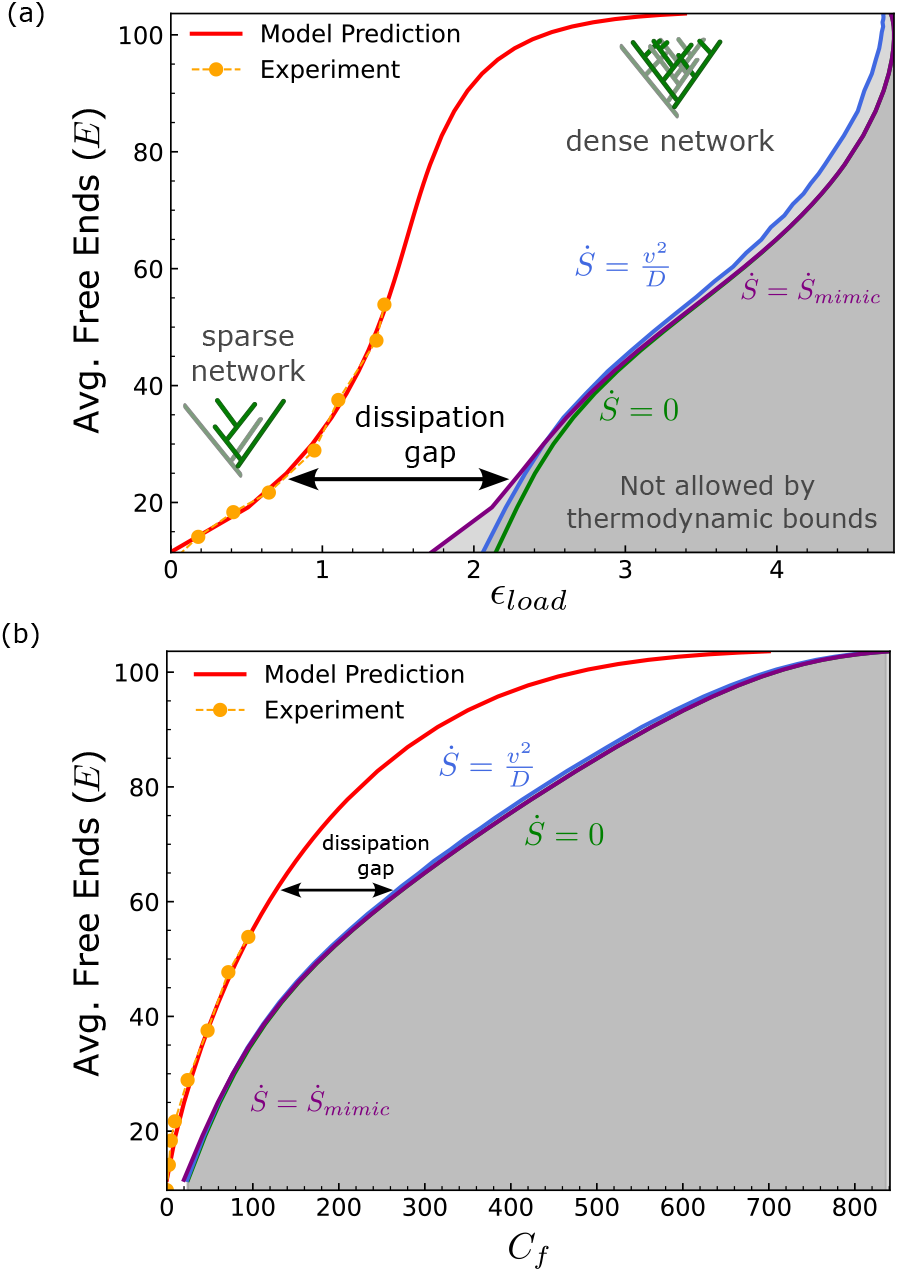
The entropy production imposes thermodynamic bounds on the nonequilibrium structure of the system, characterized by *E*. The grey shaded area indicate inaccessible configurations. (a) shows bound on *ϵ*_*load*_ from Eq. 8, while uses the bound in Eq. 9, both parametrically derived from the entropy production in Eq. 6. For a given average system density, there is some maximum load or monomer addition cost that the system has been assembling under. The distance between The model prediction and the mimic process bound reflects the amount of energy dissipated by the growing system. Yellow points in both plots represent values calculated from the experimental data of Li *et al*. [10] (Fig 1D bottom), scaled as described in Eq 13. Scaling parameters were *s* = 19.1, *β* = *−*9.5, and *κ* = .093

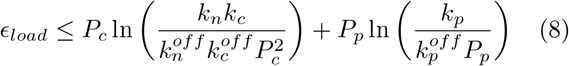

with the bound on the load in Fig. 6b as

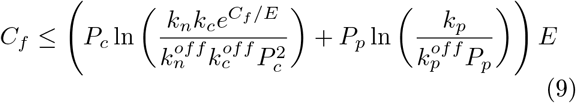

using the assumptions from Eqs. 3 and 4 we can express the monomer probabilities as

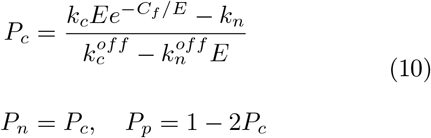

### D. A minimal non-adaptive growth process model identifies dissipative costs of maintaining adaptive response

The motion of the load due to polymerization and the maintenance of the system structure through free end density fluctuations can be viewed as linked but separable processes. The load response behavior can be recreated by a simple 1D Brownian walker. For a given system configuration defined by *E* and *P* and a load condition *C*_*f*_, we define a mimic process that takes steps of size Δ forward and back with rates

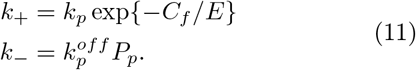

We can further define the entropy production of this mimic process as

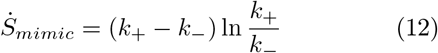

Interestingly, this entropy production rate results in bounds that are very close to the bounds discussed in the previous section (Fig. 6). Because the entropy production of this process is doing work moving the load, the rest of the entropy production of the system is dissipated to maintain the structure of the system. We call this cost a dissipation gap. In the next section we argue that this extra dissipative cost may in fact be connected to the cost associated with maintaining an adapative response in Arp2/3 networks.

### E. Adaptive properties predicted by our thermodynamic model and driven by the dissipation gap

Given that a primary function of branched actin networks is in motility through inhomogeneous networks, we next asked if we can consider our model as an adaptive system, that engages in Energy Speed Accuracy (ESA) tradeoffs between the energy dissipated by the system and the accuracy and speed of that adaptation. Applying this framework to a branched system, the adaptation problem is maintaining consistent motion (*v*_*load*_) in response to perturbation in the external environment (*C*_*f*_ ⟶ *C*_*f*_ + Δ*C*_*f*_).

The perturbation response exhibited by our model is shown in Fig. 7, where *v*_*load*_ drops immediately upon increased load, while *E* increases to the new steady state value in the slow response that adapts the system to its new conditions in an adaptation process like those described in [21]. We speculate that this trade off is fueled by the energy dissipation of the nucleation/capping processes, which can be defined as 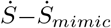. The necessity of the energy overhead needed to maintain the system is demonstrated in Fig. 7A, which demonstrates the impact of the system characteristics adapting in response to the perturbation. The mimic process uses the instantaneous values to calculate *v*_load,mimic_(*E*(*t*), *P*_*p*_(*t*), *C*_*f*_ + Δ*C*_*f*_). However, without adaptation in the system, the mimic process would predict *v*_load,mimic_(*E*_0_, *P*_*p*,0_, *C*_*f*_ + Δ*C*_*f*_), which does not maintain the desired growth, as shown in Fig 7A. For our system, we define the adaptation error as the change in *v*_*load*_ relative to the original value. The adaptation rate is extracted from fitting a logistic function to the free end recovery curve. The tradeoffs follow the same trends seen in [21], with higher energy dissipation allowing for greater accuracy, and adaptation speed upon a perturbation.

**Figure 7.**
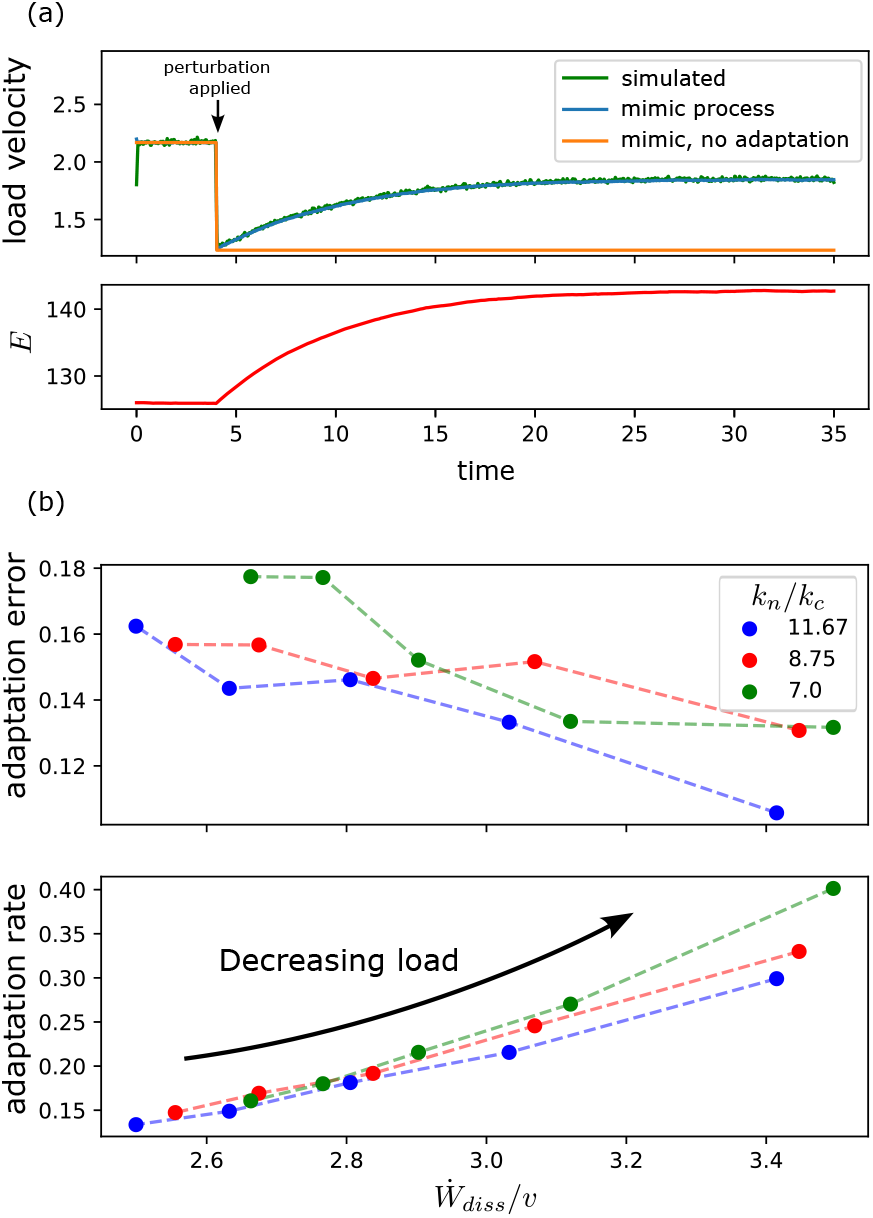
The system adapts dynamically to changes in applied load. (a) The fast response system, *v*_*load*_ drops immediately when the load is increased, while the free end density changes more slowly, acting as the mechanism of adaptation. The mimic process fully reconstructs the load velocity, even away from steady states. The effect of the shift in free ends and system composition can be seen in the yellow line, which calculates the mimic process using the unadapted *E* and *P*_*p*_ values. (b) The adaptation error (Δ*v*_*load*_*/v*_*load*0_) and rate follow the trends expected in a tradeoff with energy dissipation. Adaptation rate was determined by fitting a logistic function to the free end response curve. Energy dissipation is calculated by the entropy production per monomer addition before the perturbation.

## IV. DISCUSSION

The entropy production we calculated allows us to place thermodynamic bounds on the available structures in the system. We can also compare this to experimental data from Li *et al*. [10] (Figure 1D bottom), of normalized branched end density vs growth stress. We scaled the experimental data to our steady state solution values. The experimental data was presented in terms of the growth stress Γ and normalized branched end density *ρ*. We scaled these using fitting parameters *s, β, κ* to account for unit and normalization differences. as follows

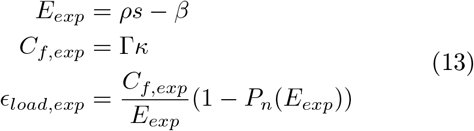

where *P*_*n*_(*E*) is calculated from Eq. 5.

The load only needed a single parameter as the zero point aligned with the simulation. This is analogous to the velocity scaling based on *c* described in Fig. 5.

The branched end densities, however, were normalized, so we could not scale them directly based off predicted rates. A second parameter was necessary to create an offset accounting for variations in the unloaded network density between our simulation and the experiment. We then optimized the parameters for the best agreement. As additional confirmation, if we estimate the rate constants using the values noted in Table S1 and use experimental reagent concentrations from [10], we get an unloaded velocity within about 8% error demonstrating further agreement with experiments.

With the qualitative agreement between the experimental and theoretical trends we can now comment on the importance of energy dissipation to maintain these non-equilibrium structures so that they can adapt to varying extracellular environments. The parametric forms inferred in Eq. 8 and 9 correspond to the quasistatic limit where no excess entropy is dissipated. These parametric forms substantially differ from those observed in experiments and in our theoretical model. This demonstrates that the energy dissipation, in the form of polymerization-depolymerization, capping, filament turnover, are all crucial to obtain the experimentally observed force morphology relations.

We also interrogate the bound on available structures using thermodynamic uncertainty relations [30]. These are a class of relations extending of the fluctuation-dissipation relation to nonequilibrium systems, describing how entropy production rates can bound the fluctuations in an evolving system. When applied to self-assembly processes, the thermodynamic uncertainty relation can be expressed as

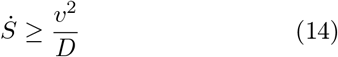

Where 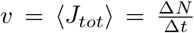, the total flux of events into the system, and 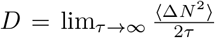. *D* captures the fluctuations in growth rate in the system, and is analogous to a diffusion coefficient [30]. This provides a tighter bound on 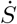 than the second law bound given by 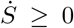. However, in this system the inclusion of the fluctuations in assembly rate did not strongly impact the thermodynamic bounds. This is likely because the system growth does not have a linear response to increasing loads. Since the parametric forms obtained in this manner differ substantially from the observed theory profiles, we can again conclude that far from equilibrium fluctuations are most likely responsible for the maintenance of the force-morphology relations of branched Apr 2/3 networks.

In summary, the bulk properties of growing branched networks under constant load can be captured in a minimal thermodynamic model. From this limited system we can determine bounds on the available structures expressed in terms of experimentally accessible quantities. Our work quantifies how dissipative processes in the background play a crucial role in allowing the system to adapt to a range of load conditions without fundamentally altering the underlying rates, determining the force-morphology-growth tradeoff curves.

This could be useful in analysis of the system, especially in interactions with heterogeneous environments where the load or resistance conditions are not fully known. This theoretical framework could be also used to analyse data from branched networks grown under unknown conditions, and potentially make inferences about the conditions the network has encountered. The model is also much more sensitive to changes in load than the underlying rates of the system making it robust to changes in background concentrations. In the future we would like to expand the thermodynamic analysis of branched networks to include responses to time-varying and heterogeneous loads such as one would encounter in cellular environments thorough interactions with membranes. We can also investigate the interactions between different nonequilibrium systems as in the interactions between assembling networks and growing membranes.

## ACKNOWLEDGMENTS

We thank Yuqing Qiu for useful discussions and Carlos Floyd for comments on the manuscript. S.V. and E.R. were supported by the National Institute of General Medical Sciences of the NIH under Award No. R35GM147400. The authors also acknowledge PHY-2317138 through the Center for Living Systems at the University of Chicago.

