## Supplemental Information for "A thermodynamic framework for nonequilibrium self-assembly and force morphology tradeoffs in branched actin networks"

### A. Effective Model

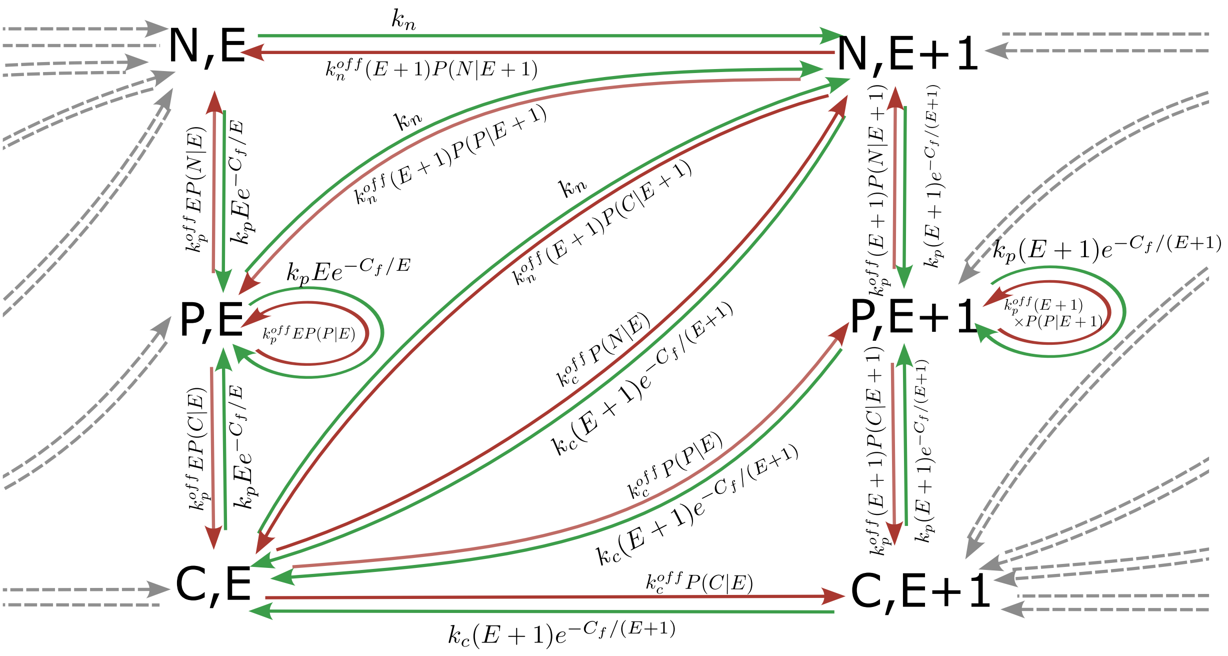


Figure S1. State transitions in the effective Markov state model. Green arrows indicates forward rates and red indicates reverse rates.

Applying the steady state approximations in Eq. 2 to the effective model in Fig. S2, we can express the master equation for the minimal model

$$
\begin{aligned}
\frac{dP_n}{dt} &= -(k_n^{off}E_{ss}(1-P_n) + k_pE_{ss}e^{-Cf/E_{ss}} + k_cE_{ss}e^{-Cf/E_{ss}})P_n + (k_n + k_p^{off}E_{ss}P_n)P_p \\
&\quad + (k_n + k_c^{off}E_{ss}P_n)P_c \\
\frac{dP_p}{dt} &= -(k_p^{off}E_{ss}(1-P_p) + k_n + k_cE_{ss}e^{-Cf/E_{ss}})P_p + (k_pE_{ss}e^{-Cf/E_{ss}} + k_p^{off}E_{ss}P_p)P_n \\
&\quad + (k_pE_{ss}e^{-Cf/E_{ss}} + k_c^{off}P_p)P_c \\
\frac{dP_c}{dt} &= -(k_c^{off}(1-P_c) + k_n + k_pE_{ss}e^{-Cf/E_{ss}})P_c + (k_cE_{ss}e^{-Cf/E_{ss}} + k_n^{off}E_{ss}P_c)P_n \\
&\quad + (k_cE_{ss}e^{-Cf/E_{ss}} + k_p^{off}E_{ss}P_c)P_p
\end{aligned}
\tag{S1}
$$

### B. Numerical comparison to experimental values

| Parameter | Value | Source |
|---|---|---|
| $k^{on}_{poly}$ | $11\mu M^{-1}s^{-1}$ | Pollard 1986 [S1] |
| $k^{off}_{poly}$ | $1s^{-1}$ | Pollard 1986 [S1] |
| $k^{on}_{cap}$ | $6.3\mu M^{-1}s^{-1}$ | Wear *et. al.* 2003 [S2] |
| $k^{off}_{cap}$ | $5 \times 10^{-4}s^{-1}$ | Wear *et. al.* 2003 [S2] |
| $k^{on}_{nuc}$ | $6.3\mu M^{-1}s^{-1}$ | Mullins *et. al.* 1998 [S3] |

Table S1. values used to estimate experimental conditions

We could not locate an experimental estimate of $k^{off}_{nuc}$, but we will assume that $k^{on}_{nuc}/k^{off}_{nuc} \approx 10^4$. Using these values and the conditions from Li *et. al.* ($5\mu M$ actin, $100nM$ CP) we can simulate the model. This resulted in an unloaded $v_{load}$ of 6.7 $\mu$M/min, which agrees with the experimental values from [S4] of 6.2 $\mu$M/min

### C. Estimation of adaptation rate

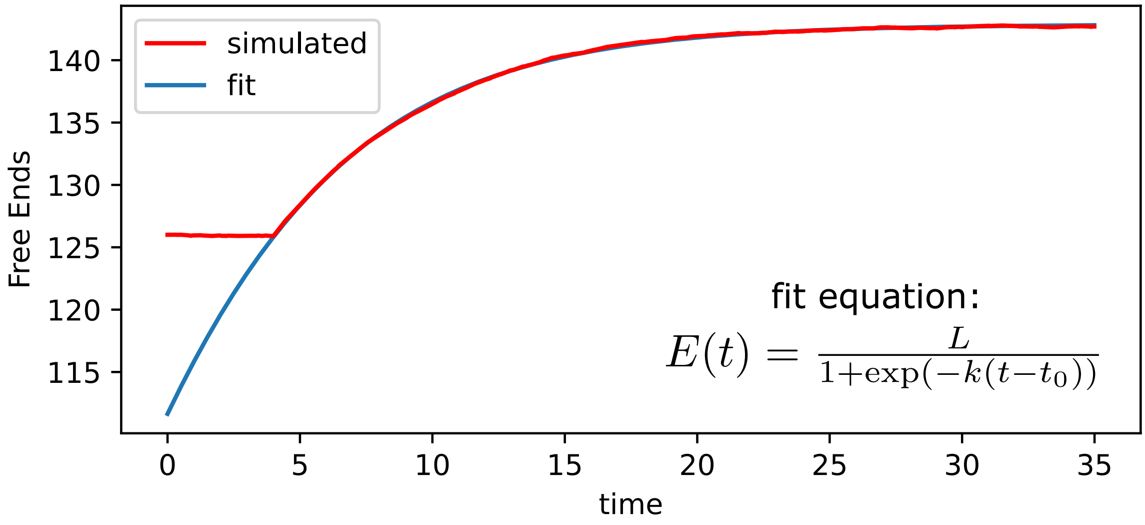


Figure S2. The adaptation rate is estimated by fitting a logistic function to the free end recovery curve. The logistic growth rate, $k$ is used as the adaptation rate. For this plot, $L = 142.9, k = 0.18, t_0 = -7.02$

[S1] T. D. Pollard, Journal of Cell Biology **103**, 2747 (1986).
[S2] M. A. Wear, A. Yamashita, K. Kim, Y. Maéda, and J. A. Cooper, Current Biology **13**, 1531 (2003).

[S3] R. D. Mullins, J. A. Heuser, and T. D. Pollard, Proceedings of the National Academy of Sciences **95**, 6181 (1998), publisher: Proceedings of the National Academy of Sciences.
[S4] T.-D. Li, P. Bieling, J. Weichsel, R. D. Mullins, and D. A. Fletcher, eLife **11**, e73145 (2022), publisher: eLife Sciences Publications, Ltd.
